# Dams have varying impacts on fish communities across latitudes: A quantitative synthesis

**DOI:** 10.1101/461145

**Authors:** Katrine Turgeon, Christian Turpin, Irene Gregory-Eaves

## Abstract

Dams are recognized to impact aquatic biodiversity and ecosystem functions, but the magnitude of effects vary across studies. By using a meta-analytical approach, we examined the effects of impoundment on fish community across three large biomes. The impacts of dams on richness and diversity differed across biomes, with significant declines in the tropics, lower amplitude but similar directional changes in temperate reservoirs, and no changes in boreal reservoirs. Our analysis also showed that non-native species increased significantly in tropical and temperate reservoirs, but not in boreal reservoirs. In contrast, temporal trajectories in fish assemblage metrics were common across regions, with all biomes showing an increase in mean trophic position and in the proportion of generalist species after impoundment. Such changes in fish assemblages may affect food web stability and merit closer study. Across the literature examined, predominant factors or mechanisms that render fish assemblages susceptible to impacts from dams were: 1) the transformation of the lotic environment into a lentic environment; 2) habitat fragmentation and 3) invasive or non-native species. Collectively our results highlight that an understanding of the regional context and a suite of metrics are needed to make robust predictions about how fish will respond to river impoundments.

## Introduction

Dams are becoming a pervasive feature of the landscape around the globe (Stickler *et al.* 2013; Grill *et al.* 2015) and hydropower has been identified by many as a clean energy source (Teodoru *et al.* 2012; Liu *et al.* 2013) that could be a major catalyst for driving the move to decarbonize our global economy (Figueres *et al.* 2017; Potvin *et al.* 2017). However, there is a clear need to identify where, and by how much dams alter the environment, particularly sensitive aquatic communities (Strayer & Dudgeon 2010). This is especially important given the unprecedent boom in dam construction in emerging economies that are mostly located in species-rich regions (Ziv *et al.* 2012; Stickler *et al.* 2013; Winemiller *et al.* 2016).

Large dams (*i.e.*, higher than 15 metres) transform large rivers into storage reservoirs, changing at least part of the ecosystem from a lotic to a lentic one (Ward & Stanford 1995; Friedl & Wüest 2002). Upstream and downstream of the dam, the alteration of the hydrological regime may generate variation in water levels and discharge far beyond natural amplitudes, with changes varying in magnitude depending on dam purpose and management (Kroger 1973; Zohary & Ostrovsky 2011). Dams can also fragment rivers by creating partial barriers to migratory organisms (Nilsson *et al.* 2005; Pelicice *et al.* 2015), or can connect aquatic ecosystems that were spatially isolated before (Gido *et al.* 2002; Gubiani *et al.* 2010). Thus, the modification of the quality, diversity, distribution and access of some key habitats should detrimentally affect some species and favor others (Stanford *et al.* 1996; Zohary & Ostrovsky 2011; Turgeon *et al.* 2018). Ultimately, dams can affect the biodiversity, ecosystem services, and the functions of the associated regulated ecosystems (Nilsson *et al.* 2005; Dudgeon *et al.* 2006; Poff *et al.* 2007; Vörösmarty *et al.* 2010).

The effects of dams on fish have been extensively studied, but divergent effects have been reported. At regional and global scales, dams can lead to fish fauna homogenization (*i.e.*, the process by which ecosystems lose their biological uniqueness; Rahel 2000; Poff *et al.* 2007; Gido *et al.* 2009; Villéger *et al.* 2011; Liermann *et al.* 2012; Vitule *et al.* 2012). At a more local scale, empirical evidence exists to show that richness and diversity decrease after impoundment, or are lower in reservoirs (Reyes-Gavilán *et al.* 1996; Pyron *et al.* 1998; Gehrke *et al.* 2002; de Mérona *et al.* 2005; Sá-Oliveira *et al.* 2015; Lima *et al.* 2016). Conversely, other studies and a recent meta-analysis (Liew *et al.* 2016) found either no change or an increase in richness and diversity after impoundment in reservoirs (Martinez *et al.* 1994a; Guenther & Spacie 2006; Irz *et al.* 2006). An increase in non-native species have been observed in several studies, suggesting that non-natives can made up the difference in total species richness in reservoirs (Martinez *et al.* 1994b; Johnson *et al.* 2008; Gido *et al.* 2009; Clavero & Hermoso 2010; Liew *et al.* 2016). Convergences among studies also suggest a general decrease in rheophilic species, an increase in limnophilic and generalist species (Bonner & Wilde 2000a; Taylor *et al.* 2001, 2014), and an increase in piscivorous fish in reservoirs (Quist *et al.* 2005; Guenther & Spacie 2006; Pelicice & Agostinho 2008; Winters & Budy 2015; Turgeon *et al.* 2018).

Earlier works have provided valuable information regarding the effects of dams on fish communities, but the divergences observed regarding fish responses to impoundment call for a global assessment that goes beyond taxonomic indices to include assemblage metrics and functional indices (Mérona and Vigouroux 2012, Mims and Olden 2013, Lima et al. 2017b). The effects might also vary across latitudes according to the inherent adaptability of fish communities to respond to the physico-chemical and biological changes brought about by dams and newly created reservoirs (Rosenberg *et al.* 1997; Gomes & Miranda 2001; Vörösmarty *et al.* 2010). Here, by using meta-analytic approach examining both taxonomic (richness, diversity and evenness) and fish assemblage metrics (number of non-native species, trophic level position and macrohabitat flow guild), across latitudes covering three large biomes, we found significant loss in richness in the tropics and relatively little change in boreal region. We also found a general change in fish assemblages toward a more generalist and predatory community, globally.

## Methods

### Literature search process

For this study, we used the guidelines and followed the checklist suggested by PRISMA (Preferred Reporting Items for Systematic Reviews and Meta-Analyses; Moher *et al.* 2009). The studies presented in this synthesis were compiled from journals indexed in Thomson ISI’s Web of Knowledge (mostly peer-reviewed articles; resulting in 668 publications) and from Google Scholar (*i.e.* peer-reviewed articles and textbooks, as well as government and industry reports, non-peer reviewed journals and conference proceedings). We searched for references including the following keywords, individually or in combination: “reservoir*”, “dam*”, “impound*”, “regulat*” but the search included “fish*” at all times. Extensive searches were performed between October 2014 and June 2017 on the references available at that time and published between 1900 and 2017. In addition, the reference lists and bibliographies of relevant sources were also scanned to find literature that was not identified through Thomson ISI’s Web of Knowledge and Google Scholar (mostly reports from the grey literature).

We then screened our database to refine our selection criteria to include only references that had quantitative data of the effects of impoundment on fish community. We thus excluded modelling and simulation exercises. A total of 67 references met our selection criteria (see Appendix S1; Table S1.1). We then classified each reference as being longitudinal (*i.e.*, one or a few reservoirs with data before and after impoundment; No. of references = 47, No. of sites = 147) or cross-sectional (*i.e.*, study presenting data on numerous reservoirs and unregulated aquatic ecosystems sampled at a single point in time; No. of references = 21, No. of sites = 37).

### Data extraction

Data were mostly extracted from tables or from datasets available in appendices and supplemental material. When data were presented in figures, they were extracted using the WebPlotDigitizer software (Rohatgi 2018). From longitudinal references, we extracted data for each reservoir and/or each sampling stations separately (*e.g.,* downstream and upstream of the dam) which sometimes resulted in more than one study site per reference (147 studies from 47 references; Table S1.1). The analysis was performed at the study site level for longitudinal studies. From each longitudinal study, in addition to fish data over time, we consistently recorded: 1) the archetype of the reservoir visually assessed (lake-shaped, dendritic and canyon; Appendix S2, Fig. S2.1), 2) the geographic location (longitude and latitude), 3) the freshwater ecoregion (Abell *et al.* 2008; http://www.feow.org/), 4) the location of the sampling station (downstream, upstream of the dam or upstream of the reservoir), 5) the distance from the dam, 6) the duration of the study, 7) the area of the reservoir at full pool, 8) the flooded terrestrial area, 9) the catchment area (or watershed area), 10) the main reservoir usage (hydroelectricity, water storage, irrigation, flood control, multi-purpose) and 11) the main mechanisms reported by the authors to be responsible of the observed change in diversity or fish assemblages (Table S1.1). We aimed to collect a similar set of data from the 47 cross-sectional studies, including: 1) the geographic location (longitude and latitude), 2) the freshwater ecoregion(s), 3) the location of the sampling stations, 4) the area of the reservoir and reference lakes when available, 5) key dates including the date at which the reservoir was created, the date at which the river or lake was impounded and the date at which the reservoir reached its full pool, and 6) the main mechanisms responsible of the observed change in assemblages (Table S1.1). In cases where these data were not presented in the original references, we attempted to find these data from alternative sources (*e.g.*, Google Earth or other published studies focused on the same target ecosystem). See Appendix S2 (Fig. S2.2) for a graphical summary of the dataset.

#### Calculation of the taxonomic metrics

##### Richness

Richness values (*i.e.*, the number of fish species) were provided in all studies. We also used *ln* values of richness, where trends can be interpreted as an estimation of proportional changes per biome when values lie between −1 and 1. *Diversity* – Values of diversity were directly provided in only three studies. However, many references had data on relative abundance of the species in the community (24/47 studies for longitudinal and 17/37 studies in cross-sectional studies). We used these relative abundance data to calculate diversity, evenness, the mean trophic level position and macrohabitat flow guild. We calculated diversity by using the Shannon’s H’ diversity index 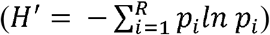, The Shannon’s H’ takes evenness and species richness into account and quantifies the uncertainty in predicting the species identity of an individual that is taken at random from the dataset and where *p_i_* is the proportion of individuals belonging to the *i*^th^ species in the dataset. *Evenness* – We calculated evenness by using the Pielou’s J’ Evenness index. Pielou’s 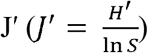 ranges from near 0 (indicating pronounced dominance) to near 1 (indicating an almost equal abundance of all species) and H’ is the Shannon’s H’ diversity index where S is the total number of species.

#### Calculation of the change in fish assemblages’ metrics

##### Non-native species

When provided, we extracted the number of non-native species observed. In this contribution, a non-native species consisted of a species that is introduced beyond its native range as a direct (*e.g.*, stocking angling, bait fish) or indirect result of human action (elimination of the barrier that connects adjacent aquatic ecosystems through “natural” dispersal; Jeschke *et al.* 2014). *Trophic level position -* We extracted the mean trophic level position for each species from FishBase (Froese & Pauly 2015), and we calculated a mean trophic level position metric using: 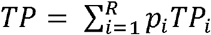, where *p_i_* is the proportion of individuals belonging to the *i*^th^ species and *TP_i_* is the average reported trophic level position for species *i*. *Macrohabitat flow guild* – We first categorized fish species based on their macrohabitat flow guild (generalist, fluvial facultative or fluvial specialist) by using FishBase (Froese & Pauly 2015) and other articles and books (Scott & Crossman 1973; Travnichek & Maceina 1994; Quinn & Kwak 2003; Guenther & Spacie 2006; Baumgartner *et al.* 2014; Buckmeier *et al.* 2014; Lima *et al.* 2017a). Generalists species were coded 1, whereas fluvial facultative taxa were coded 0.5 and fluvial specialists coded 0. We then used this formula to generate an index of macrohabitat flow guild, 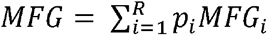, where *p_i_* is the proportion of individuals belonging to the *i*^th^ species and *MFG_i_* is the macrohabitat guild for species *i* (the MFG metrics varies from 0 to 1).

### Data analysis

We ran separate analyses for longitudinal (data before and after impoundment) and cross-sectional datasets (data comparing reservoirs and unregulated aquatic ecosystems). For longitudinal studies, our main goal was to extract trends regarding the impacts of impoundment over time on fish metric across studies (*i.e.*, metrics ∼ time since impoundment). Conventional meta-analyses rely on the assumption that sampling distributions have known conditional variances (*i.e.*, weight assigned to both the variances and sample sizes from original studies) and that effect size estimates from different studies are independent (Borenstein *et al.* 2009; Hedges *et al.* 2010; Gurevitch *et al.* 2018). In our synthesis, we could not satisfy the assumption of known conditional variance because for many of our longitudinal studies (39%), sample size variance was unknown or could not be calculated. We were also interested to use sampling station as our statistical unit (called “studies”) and thus our studies were not independent (some stations came from the same reservoir). Because of these limitations, we ran Linear Mixed Effects Models (LMM; lmer function in the lme4 library v.1.1-18-1, Bates *et al.* 2018) that were weighted by the number of observation in the time series (and assumes that sample size is inversely related with variance). In addition, the application of LMM allowed us to increase our power (Hillebrand & Cardinale 2010) and to add a complex structure of covariates in the fixed effects that cannot be easily implemented in most meta-analysis packages. All analyses were performed in R (v. 3.3.2; R Core Team 2017).

We ran three sets of models to fully understand the observed patterns because we were really interested in how taxonomic and fish assemblage metrics varied across biomes. We first ran a model for the combined dataset (where all biomes were combined). Secondly, we ran a model comparing biomes among each other and used the interaction between time and biome in the LMM (covariates). Finally, we ran separate models per biome to examine how richness changes over time per biome, without drawing comparisons among biomes. For the three sets of models, we used (years | study_ID/Reservoir_ID) as random factors, controlling for the effect of time per study, and where each study was nested in its reservoir (to control for the spatial non-independence of the studies; similar approaches have been used in Liao *et al.* 2007; Rey Benayas *et al.* 2009; Vilà *et al.* 2011). The mean effect size for the fixed effects was estimated with Restricted Maximum Likelihood and calculated with Kenward-Roger approximation to approximate degrees of freedom in mixed effect models (Kenward & Roger 1997) by using the sjPlot package in R (v. 2.6.0; Lüdecke & Schwemmer 2018).

For the cross-sectional studies, since both variances and sample sizes were available, we ran conventional weighted meta-analyses. We weighted effect size estimates by their inverse variance weights, such that studies with higher sample sizes were given more weight by using (*i.e.*, weights = ((1/SD)*N), following Borenstein *et al.* 2009; Hedges *et al.* 2010). We assessed differences in the overall effect size (*e.g.,* if the mean of each metrics differs between regulated or unregulated ecosystems) by using the Standardized Mean Difference (SMD). For each biodiversity metric considered in the cross-sectional studies, we also ran three sets of models (one for the combined effect, one with the interaction term (biome*effect), and separate models per biomes). Unfortunately, not enough information was provided for non-native species in cross-sectional studies.

We used regression trees (rpart package, v. 4.1-10, Therneau *et al.* 2018) to explore and explain the variability observed and residual variance in random effects values (RE) of the mixed models based on reservoir or sampling station characteristics (when relevant, see Table S1.1 and Fig. S2.2). Regression trees were pruned by minimizing the cross-validated error to avoid overfitting (De’ath & Fabricius 2000).

### Publication bias

We explored the possibility of publication bias by using funnel plots (Appendix S5), which allow for a visual assessment of whether studies with small effect sizes are missing from the distribution of all effect sizes (*i.e.*, asymmetry). We also ran Spearman rank correlations to examine the relationship between the standardized effect size and the sample size across studies, and the relationship between the standardized effect size and the duration of the studies for longitudinal studies (Rosenberg *et al.* 2000). A significant correlation would indicate a publication bias whereby larger effect sizes are more likely to be published than smaller effect sizes, when sample size is small or duration of the study is short.

## Results

### How do impoundments affect fish biodiversity and assemblages? Longitudinal studies

#### Taxonomic metrics (Richness, Diversity and Evenness)

Richness and diversity decreased significantly over time when all studies and regions were combined, but this pattern is observed only in temperate and tropical regions when biomes were modeled separately (Fig. 1 a, c). Richness and diversity decreased at a much faster rate in tropical reservoirs when compared to boreal and temperate reservoirs (Time: B vs. TR, boreal used as the contrast in the model; Fig. 1 a, c). Using *ln* richness, we still observed a decrease over time in the combined dataset, as well as in temperate and tropical regions (Fig. 1 b), but the proportional rate of change in richness over time is comparable across biomes (interaction terms between Time and Biomes are not significant; Fig. 1 b). Evenness did not significantly decrease over time for the combined dataset. However, we did find that evenness declined significantly in tropical and temperate regions when biome-specific models were performed (Fig. 1 d). Compared to boreal reservoirs, evenness was lower and decreased faster in temperate regions (Fig. 1 d).

**Figure 1.**
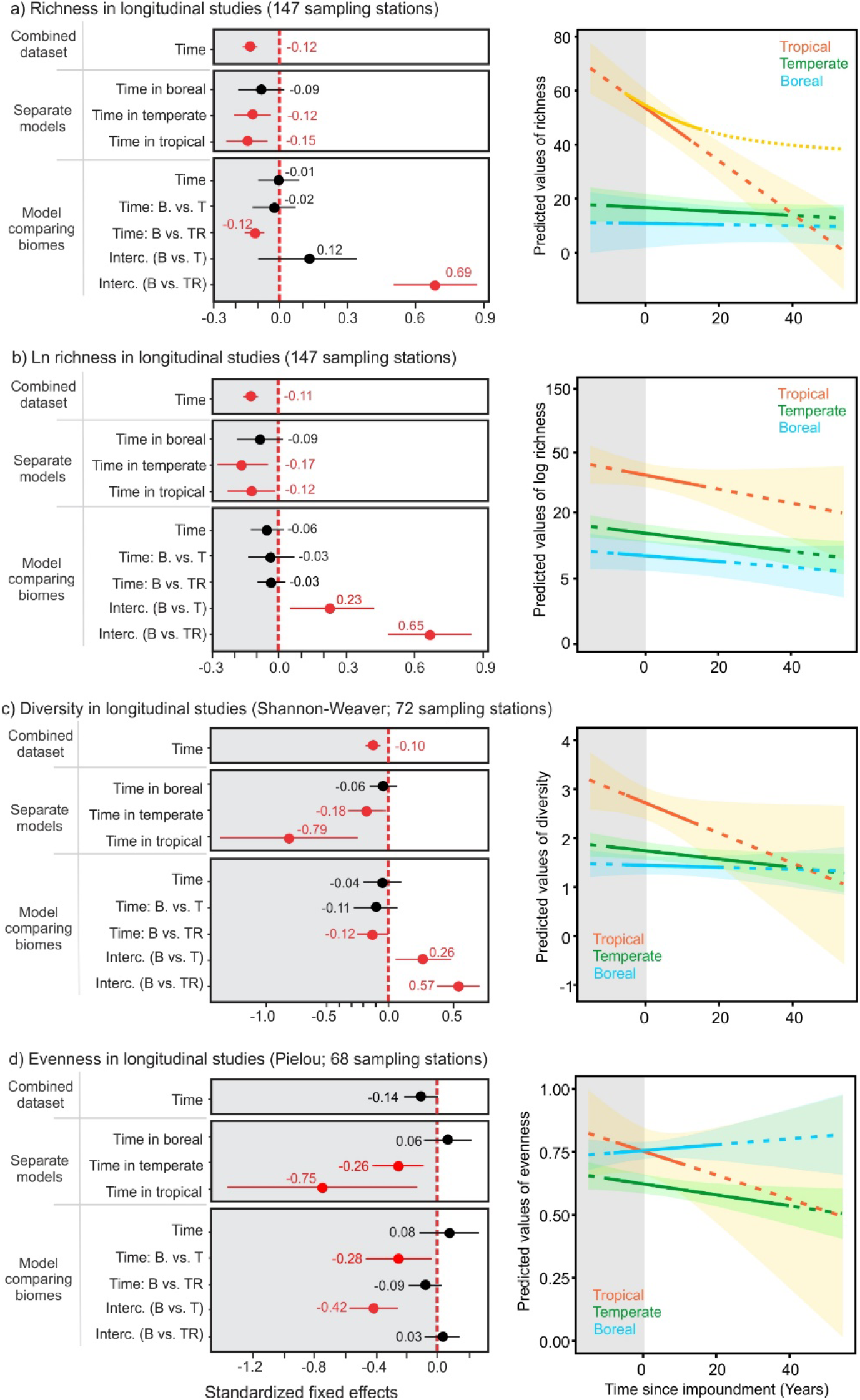
Standardized effect size in longitudinal studies for taxonomic metrics. *Left panels*: Standardized fixed effects coefficients and standard error from the mixed effects models comparing the temporal trends in a) richness, b) *ln* richness, c) diversity, and d) evenness across biomes. For each taxonomic metric, we showed the effect of time since impoundment on the metrics for the combined dataset, for each biome modelled separately, and we also compared biomes in the model by using an interaction between biome and time (Time: B vs. T and Time: B vs. TR, using boreal (B) as the contrast). The 95% CI were evaluated with the Kenward Roger approximation. Coefficients in red represent significant patterns for a given fixed effect. *Right panels:* Models predictions from the model comparing the effect of time since impoundment on taxonomic metrics across biomes (model with the interaction). The solid lines represent the predictions where we have actual data and the dotted lines represent model extrapolation. In a), the non-linear curve in the tropics is a more plausible pattern but cannot be modeled.

#### Species assemblage metrics (Non-native, Trophic level position and Macrohabitat flow guild)

Across all analyses of species assemblage metrics, we detected different results when the combined datasets (studies across all biomes) were contrasted with the regionally-specific analyses. For example, no non-native species were observed in any of the boreal reservoirs following impoundment, but some invaded tropical and temperate reservoirs (Fig. 2 a). The number of non-native species increased for the combined dataset, but this was driven by tropical and temperate regions (Fig. 2 a). We found that the rate of change in non-native species increased much faster in tropical reservoirs than in temperate reservoirs (Fig. 2 a), and this was still true when the proportional rate of increase in non-native species was considered (Fig. 2 b). The average trophic level position increased following impoundment for the combined dataset, as well as in boreal reservoirs, but not in temperate and tropical reservoirs (Fig. 2 c). The mean trophic level position was lower in temperate and tropical regions when compared to the boreal, and the rate of change did not differ across regions (Fig. 2 c). We observed an increase in generalist species over time in boreal and tropical regions as well as in the combined dataset (but did not in temperate region alone; Fig. 2 d). Compared to boreal reservoirs, temperate and tropical reservoirs had a higher proportion of rheophilic species originally (Fig. 2 d) and the rate of change in the proportion of generalist species increased more slowly in temperate than in tropical regions (Fig. 2 f).

**Figure 2.**
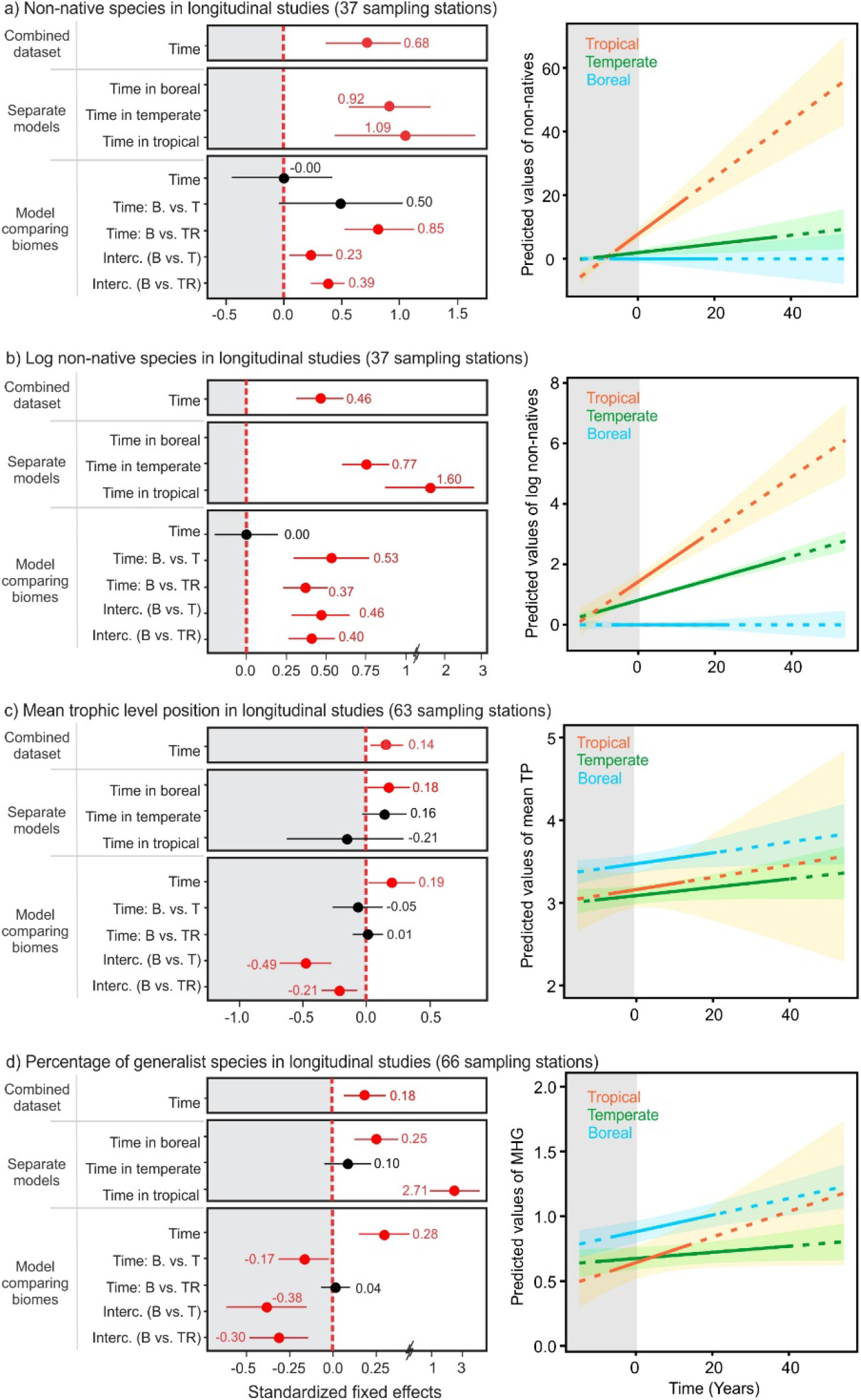
Standardized effect size in longitudinal studies for taxonomic metrics. *Left panels*: Standardized fixed effects coefficients and standard error from the mixed effects models comparing the temporal trends in a) number of non-native species, b) *ln* number of non-native species, c) mean trophic level position, and d) percentage of generalist species across biomes. For each taxonomic metric, we showed the effect of time since impoundment on the metrics for the combined dataset, for each biome modelled separately, and we also compared biomes in the model by using an interaction between biome and time (Time: B vs. T and Time: B vs. TR, using boreal (B) as the contrast). The 95% CI were evaluated with the Kenward Roger approximation. Coefficients in red represent significant patterns for a given fixed effect. *Right panels:* Models predictions from the model comparing the effect of time since impoundment on taxonomic metrics across biomes (model with the interaction). The solid lines represent the predictions where we have actual data and the dotted lines represent model extrapolation.

#### Study specific effects (random effect values; RE)

The examination of the study specific effects as random effect values (RE; forest plot) from the mixed effects models showed a much higher variability in richness in the tropics than in temperate and boreal regions and showed that residual variances across studies (at the sampling station level) were comparable within a given reservoir (Appendix S3; Figs. S3.3, and S3.5). Using a regression tree on the RE of the model built with tropical data only, we found that variation in RE was significantly associated with the catchment area of the reservoirs (Proportional reduction in error (PRE) = 54.6% of the variation explained) and the duration of the study (PRE = 9.3%; Fig. 4). Reservoirs with large catchment area showed a tendency to experience a higher loss of species relative to the mean loss of richness in this region, whereas young reservoirs with smaller catchment area experienced a lower loss of species, and sometimes even an increase in richness (Fig. 4). RE values for diversity and evenness did not show this amount of variability (Fig. S3.6). The examination of the RE values for the number of non-native species showed some variability in the tropics but very little in boreal and temperate regions (Fig. S3.7). Variability across studies was not significant in the other species assemblage metrics (Figs. S3.7 and S3.8).

### How do impoundments affect fish biodiversity and assemblages? Cross-sectional studies

#### Diversity metrics (Richness, Diversity and Evenness)

Across the different diversity metrics, we could obtain a substantially larger number of observations for richness, relative to diversity and evenness. Based on our analyses of the cross-sectional studies, richness did not differ between regulated (reservoirs and regulated rivers and streams) and unregulated aquatic ecosystems when all studies and regions were combined. However, when using separate models, we found higher richness in regulated ecosystems relative to unregulated ecosystems in temperate region (Fig. 3 a). Diversity was higher in regulated ecosystems compared to in unregulated ones when tropical and temperate ecosystems were considered separately (Fig. 3 b). The difference in diversity between regulated and unregulated ecosystems was greater in temperate region when compared to boreal and tropical regions (Fig. 3 b). Evenness did not differ between regulated and unregulated aquatic ecosystems for the combined dataset but was higher in regulated tropical and temperate ecosystems when each biome was considered separately (Fig. 3 c). The examination of the RE values for taxonomic metrics did not show significant heterogeneity across individual studies (Appendix S3).

**Figure 3.**
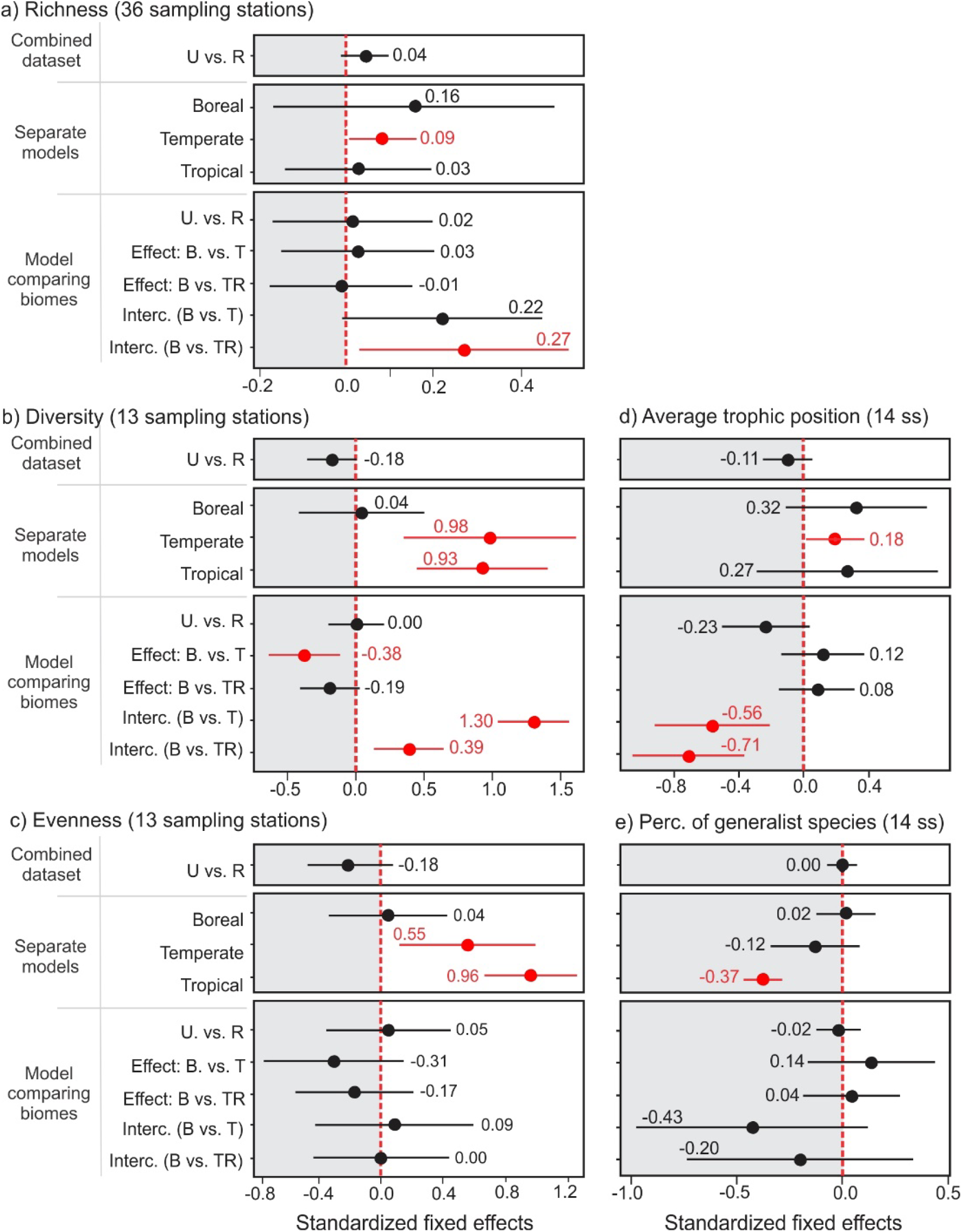
Standardized effect size in cross-sectional studies for taxonomic and fish assemblage metrics. Standardized fixed effects coefficients and standard error from the mixed effects models comparing the temporal trends in a) richness, b) diversity, c) evenness, d) mean trophic level position, and e) percentage of generalist species across biomes. Positive deviations in standardized fixed effects represent cases where the regulated ecosystems had higher values relative to the unregulated ecosystems. For each metric, we showed the difference between regulated (R) and unregulated ecosystems (U) on the metrics for the combined dataset, for each biome modelled separately, and we also compared biomes in the model by using an interaction between biome and effect (Effect: B vs. T and Effect: B vs. TR, using boreal (B) as the contrast). The 95% CI were evaluated with the Kenward Roger approximation. Coefficients in red represent significant patterns for a given fixed effect. Coefficients in red represent significant patterns for a given fixed effect.

#### Assemblages metrics (Non-native species, Trophic level and Macrohabitat flow guild)

Regarding species assemblage metrics in cross-sectional studies, few patterns were significant. The average trophic level position did not differ between regulated and unregulated aquatic ecosystems for the combined dataset, nor did it for the boreal or tropical regions when examined separately, but was higher in regulated ecosystems in temperate region (Fig. 3 d). The percentage of generalist species did not differ between regulated and unregulated ecosystems for the combined dataset, for boreal and temperate regions, but was lower in tropical regulated ecosystems (Fig. 3 e). We did not have enough data to examine if the number of non-native species differ between regulated and unregulated ecosystems across all biomes. The examination of the RE values for species assemblage metrics also did not show significant heterogeneity across individual studies (Appendix S3).

## Discussion

### Gradient of impacts on biodiversity across latitudes

The impacts of dams on fish biodiversity followed a clear gradient across latitudes, from a general lack of apparent changes in boreal regions to substantial ones in the tropics. A previous meta-analysis by Liew *et al.* (2016) suggested that dams have similar effect across regions, but their analyses did not consider the boreal region. In addition, we report on a substantially larger pool of information, representing 60% increase in number of references considered by Liew *et al.* (2016). As such, the gradient of effects we report on here clearly underscores the need for an understanding of regional fish assemblages, and the context of stressors when evaluating the impacts of damming rivers on fish biodiversity.

Fish from tropical rivers and temperate prairie streams have evolved in fluvial ecosystems and most lack the morphological, behavioural and reproductive traits, as well as plasticity needed to successfully occupy the new lentic habitats created upstream of the dam (Gomes & Miranda 2001; Dodds *et al.* 2004; Agostinho *et al.* 2008; Durham & Wilde 2011). Such a lack of traits and plasticity can partly explain the decrease in richness observed over time in longitudinal studies in tropical and temperate ecosystems. On the other hand, boreal reservoirs from this synthesis have minimal anthropogenic impacts other than dams due to their remote locations, and have no reports of non-native species (Sutela & Vehanen 2008; Turgeon *et al.* 2018). Large lakes are also much more common in the boreal region than in temperate and tropical regions (Verpoorter *et al.* 2014; Messager *et al.* 2016), and fish have been colonized boreal aquatic ecosystems from refugia after glaciers began retreating about 15 000 years ago (Schluter & Rambaut 1996; Griffiths 2006). For these reasons, boreal freshwaters fish fauna is depauperate and characterized by large body size species that are generally able dispersers and ecologically-tolerant species (Dynesius & Jansson 2000; Griffiths 2006; Lévêque *et al.* 2008). Collectively, these characteristics make boreal fish communities potentially quite resilient to river impoundment.

Given the differences in the length of available time series across biomes and the substantial heterogeneity observed across reservoirs within the tropics, it is important to scrutinize the data before drawing generalizations about the sensitivity of fish richness and diversity in impounded tropical systems. Tropical reservoirs are much younger than temperate and boreal reservoirs, and therefore time series available for the tropics are shorter (*i.e.*, on average 6 years, as opposed to 18 or 19 years as found with boreal regions and temperate regions, respectively; Fig. S2.2). For completeness, we truncated the time series in temperate and boreal regions to only keep richness data spanning up to 5 y post- impoundment, and re-analysed the data (Appendix S4, Fig. S4.10). Even with comparable study periods (5y), richness still decreased faster in the tropics (Fig. S4.10). We must still be careful with predictions that extend beyond 10 years in duration in the tropics because they could well be overestimating loss in richness (Fig. 2 a; dashed line). An alternative and more plausible trajectory would be a decreasing non-linear curve that stabilizes at some point (Fig. 2 a; saturating curve illustrated), but the short time series did not allow us to test for non-linear patterns over time. Furthermore, we observed significant heterogeneity across studies in the tropics when compared to temperate and boreal regions (Figs. S3.3 and S3.5). This variability across studies was significantly associated with the size of the catchment area and the duration of the study (Fig 4). A higher decrease in richness relative to the mean loss of species was observed in reservoirs located in large catchment area. Rivers in larger catchment areas usually have higher richness (Welcomme 2000; in this study: LMM, estimate ± SE = 0.242 ± 0.078, P = 0.003, R^2^ = 0.17), and thus had a higher potential to lose species. A lower loss in richness relative to the mean loss of species or an increase in richness was observed mostly in short duration time series and can partly be explained by the short term and rapid increase in non-native species (supported by this meta-analysis) that were better adapted to the newly created lentic habitats (Rahel 2002; Clavero & Hermoso 2010; Vitule *et al.* 2012) and may result in biotic homogenisation at larger scales (Poff *et al.* 2007; Gido *et al.* 2009; Vitule *et al.* 2012). These non-native species can come from newly connected drainages by the flooding of natural barriers (direct effect; Júlio *et al.* 2009; Clavero & Hermoso 2010; Vitule *et al.* 2012), or by intentional or unintentional species introduction (indirect effect through propagule pressure; Johnson *et al.* 2008; Pelicice & Agostinho 2008). However, this increase in richness in the tropics is suggested to be transient because some studies demonstrated a rise and fall in richness (humped-shaped non-linear pattern) after impoundment in the tropics (Agostinho *et al.* 1994; Lima *et al.* 2016), stressing for the need of longer time series.

**Figure 4.**
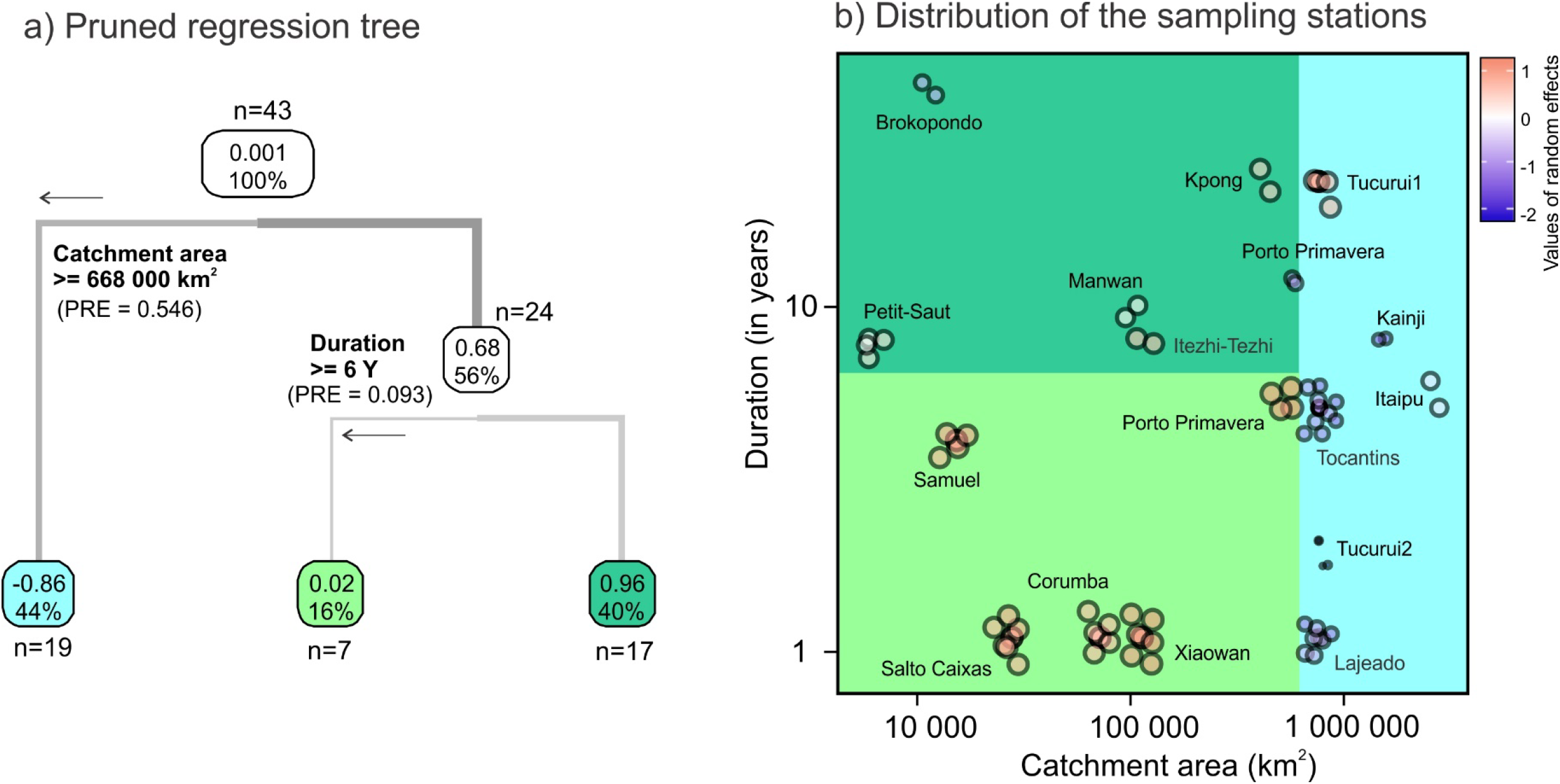
Regression tree predicting the heterogeneity in random effect values (RE) from the mixed effect model examining the effect of time since impoundment on richness in the tropics. a) plot of the pruned regression tree showing the mean value and proportion of the dataset in boxes at each step of the tree, and b) distribution of the sampling stations and reservoirs in the catchment area and duration of the studies space. The size and color of the circle represent the RE values of the model (i.e., variation relative to the mean loss of richness in this region). Red means a lower loss in richness relative to the mean loss of richness in this region and an increase in richness in some cases, blue means a higher decrease in richness relative to the mean loss of richness in this region. The different shades area in panel b correspond to data range covered by the three final nodes in the pruned regression tree.

### Impacts on the food web: Less rheophilic, more non-native and predatory species

Our meta-analytic approach suggested a global decrease in rheophilic species (more pronounced in tropical region), an increase in non-native (except in the boreal region) and an increase in mean trophic level position (Fig. 2). A decrease in rheophilic species was expected following the transformation of a lotic to a lentic ecosystem (Gomes & Miranda 2001; Agostinho *et al.* 2008) due to strong selective pressures in these newly created lentic habitats that should favor generalists over rheophilic and fluvial specialist species (Li *et al.* 2013). Interestingly, the strength of trophic interactions and the observed increase in predatory fish in reservoirs can also contribute to the decrease of rheophilic species. Increased predator densities have been suggested to reduce migration success of small-bodied stream fishes (Matthews & Marsh-Matthews, 2007; Franssen, 2012).

The general increase in the trophic level position can be due to an increase in predatory fish (higher trophic level position), a decrease in benthivorous and planktivorous fish (lower trophic position) or to both mechanisms. Because reservoirs are frequently larger and more accessible to humans relative to natural lakes, they attract significant numbers of recreational fisherman. Likewise, reservoirs have been subject to intense fish stocking and species introduction, mainly for piscivores and sport/game fish species (Pelicice & Agostinho 2008). Water drawdown in reservoirs can also favor piscivores by concentrating prey fish (Hulsey 1956; Ploskey 1986; Nordhaus 1989; Sutela & Vehanen 2008), which can increase the feeding activity and growth of young and adult piscivores (Heman *et al.* 1969; Zweiacker *et al.* 1972; Johnson & Andrews 1973; Heisey *et al.* 1980; Herrington *et al.* 2005). Moreover, the trophic surge following impoundment can also benefit predators by the boom of productivity during and shortly after impoundment, but this effect might be transient. Lastly, the cannibalism observed in many large predators in reservoirs might keep reservoirs in a predator-dominated state (McCauley *et al.* 2018) and might confer some stability to the food web (Claessen *et al.* 2004; McCann 2011).

These changes in species assemblages, and how they can impact the structure and the stability of food webs in reservoirs deserve closer investigation, especially in the tropics where alterations to species-rich food web are greater, on-going, and not well-understood (Layman *et al.* 2005; Rooney *et al.* 2006). Reservoirs seem to have longer food chains (Hoeinghaus *et al.* 2008; Mercado□Silva *et al.* 2009) and more “weblike” interactions, especially in the presence of omnivory (Stein *et al.* 1995). The potential impacts of dams on food web stability call for a better integration of taxonomic, functional and life history trait responses to impoundment at a global scale (Mérona & Vigouroux 2012; Mims & Olden 2013; Lima *et al.* 2017b) because their relative importance can change across latitude and in a spatio-temporal context.

### Mechanistic understanding of the effects of impoundment on fish assemblages

Several hypotheses regarding the mechanisms responsible for the change in biodiversity and fish assemblages following impoundment have been suggested. As a first exploratory step to develop a general mechanistic understanding, we extracted the main mechanisms reported by the authors from our 67 references (Fig. 5; Table S1.1). We then classified the 11 identified mechanisms into three categories: 1) alteration of the hydrological regime, 2) impacts on connectivity and fish movement and 3) change in food web and trophic interactions (Fig. 5).

**Figure 5.**
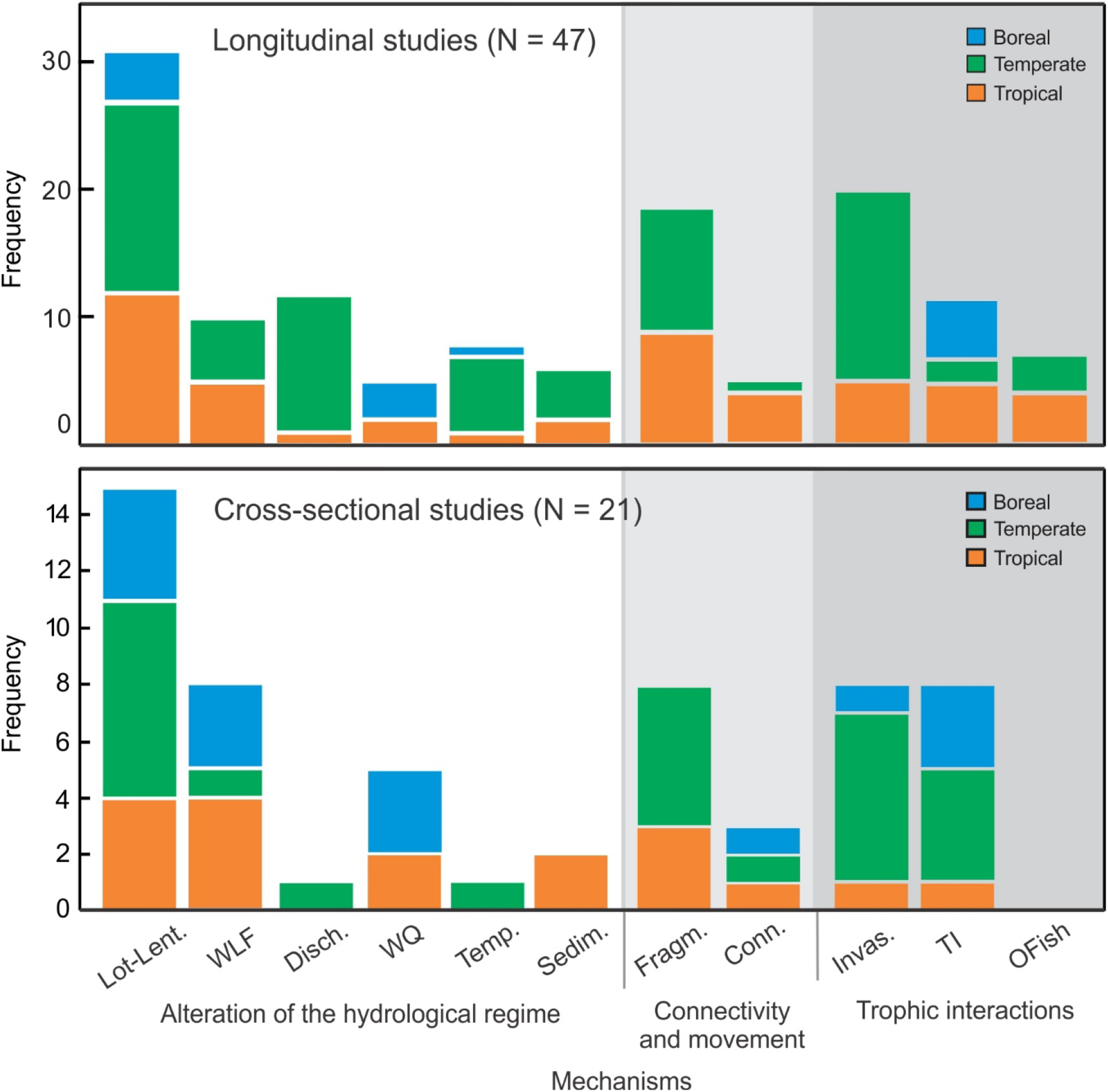
Summary of the main mechanisms affecting fish biodiversity in a) longitudinal and b) cross-sectional studies. Frequency distribution of the main mechanisms reported by the authors (in the abstract and/or conclusions) and potentially responsible for the change in fish assemblages observed in the 67 references. The mechanisms were classified into three main classes: Alteration of the hydrological regime, alteration of the connectivity and fish movement, and impacts on the trophic interactions. Lot-Lent. = change from a lotic to lentic condition, WLF = water level fluctuation in the reservoir, Disch. = change in discharge downstream of the dam, WQ = change in water quality excluding temperature, Temp. = change in temperature upstream and downstream of the dam, Sedim. = change in sedimentation regime, Fragm. = dams fragment river dynamics and can create a barrier to movement, Conn. = increased connectivity of the drainage basins, Invas. = increase in the number of invasive species, TI = change in the strength of the trophic interactions (predation and competition), OFish = overfishing of some species.

The alteration of the hydrological regime can affect fish communities by shifting the ecosystem from a lotic to a lentic one, through changes in discharge and water levels, and by changing water quality, temperature and sedimentation regimes (Fig. 5). The transformation of the lotic environment into a lentic environment was the most commonly cited mechanism (69% of the studies; Fig. 5). The new lentic conditions upstream of the dam and a change in discharge downstream can adversely affect fluvial specialists and large-river species (Winston *et al.* 1991; Bonner & Wilde 2000b; Franssen & Tobler 2013; Taylor *et al.* 2014); this mechanism was clearly illustrated in our meta-analysis by the general decrease in rheophilic species and an increase in generalists. Water levels fluctuations and winter drawdown can affect fish that depend on the littoral zone through modification of their feeding, growth and reproduction (freezing of eggs and larvae, lost of spawning substrate; June 1970; Gafny *et al.* 1992; Kahl *et al.* 2008; Probst *et al.* 2009) and also indirectly through changes in prey availability and quality (Paller 1997; Furey *et al.* 2006; Aroviita & Hämäläinen 2008; Zohary & Ostrovsky 2011; Stoll 2013). Information on the proportion of benthophages or species inhabiting the littoral zone would help inform this later mechanism.

The modification of the riverscape connectivity by dams can also alter fish assemblages by limiting the movement of migratory species, by affecting metapopulation dynamics, or by facilitating invasions by connecting aquatic ecosystems (Dynesius & Nilsson 1994; Fullerton *et al.* 2010). The fragmentation of rivers through the construction of barriers to migration was another mechanism commonly cited (39% of the studies, Fig. 5). Populations isolated in upstream areas by dams can be subject to extirpation when reproductive failure or high mortality cannot be counterbalanced by recolonization from downstream sources (Winston *et al.* 1991). On the other hand, some authors have observed increased colonization of non-native species in impounded streams (Havel *et al.* 2005; Johnson *et al.* 2008). To capture this mechanism in future meta-analyses, the proportion of species undergoing migration (anadromous, potamodromous) needs to be reported more frequently.

In addition to a higher susceptibility to propagule pressure, reservoirs are particularly vulnerable to successful establishment of non-native species (40% of the studies, Fig. 5) because they are in a perturbed state after impoundment compared to natural lakes (Thornton *et al.* 1990; Pringle *et al.* 2000; Davis 2003; Didham *et al.* 2007). Several studies have found an increase in non-native species after impoundment, and often these taxa are piscivorous species that become quite abundant post-impoundment (Martinez *et al.* 1994b; Quist *et al.* 2005; Guenther & Spacie 2006; Johnson *et al.* 2008; Gido *et al.* 2009; Clavero & Hermoso 2010; Franssen & Tobler 2013). When introduced, they compete with, and can prey upon native species (Li et al. 1987, Minckley et al. 1991). Basses are well known to homogenize fish assemblages by eliminating small-bodied prey species (Jackson 2002) and are very often introduced in temperate reservoirs. Quist *et al.* (2005) found that the Great Plains river fish assemblage switched from a catostomids and cyprinids (*i.e.* river specialists) dominated system prior to impoundment to an non-native species assemblage, mainly dominated by piscivores (*e.g.*, smallmouth bass, walleye, yellow perch and brown trout).

The above-mentioned mechanisms are mostly based on authors opinions and not on strong quantitative evidence because replication in longitudinal studies is rare. The dominant mechanisms can also differ according to the location and scale (biomes, location of the sampling station), may be dynamic over time (*i.e.,* differ among the filling phase vs. shortly after or many years after impoundment) and can be influenced by the particularities of reservoir management and confounding factors (*i.e.*, stocking, fishing). This summary also enlightens the importance of moving toward a trait-based approach to get a mechanistic understanding of the effects of impoundment on fish communities, which needs to be more thoroughly reported in original research studies.

### Limitations and publication bias

The main limitations and/or biases in our synthesis that could affect the interpretation and strength of evidence are: 1) publication bias, 2) variation in fishing effort and gears, 3) variation in the duration of the studies, 4) assumption of a linear relationship between time and richness, 5) calculation of the trophic position in a changing habitat, 6) defining an adequate reference ecosystem for a reservoir, and 7) the difference in ecosystems size. We addressed the issue of publication bias with the visual inspection of funnel plots and Spearman rank correlation examining the effect size in relation to the sample size and duration of the study (Appendix S5). Funnel plots show an absence of a clear sampling/publication bias in most cases, but we found few significant Spearman r values suggesting a bias toward publishing large effect sizes when sample size is small or study duration is short (Appendix S5). Second, the effort and the fishing gears used varied across studies, but also among years in some studies. Roughly 41% of the studies did not have similar effort across years (See Table S1.1) – sometimes using different fishing gears - and only 23% of the studies reported rarefied richness (*i.e.*, controlling for the number of samples). Most studies used gill nets, resulting in an underestimation of small littoral and pelagic species. Third and fourth, the duration of the study also varied among studies and was much shorter in the tropics. The consequences and implications of these limitations were discussed earlier. Given the predominance of shorter time series, particularly in the tropics, we assumed a linear relationship between time and richness; with longer time series, nonlinear modeling would be worth exploring. Fifth, we assumed that the change in habitat brought about by dam would not change the trophic level position for a given species. However, some studies have demonstrated that, in altered habitats or those invaded with non-native species, the trophic level position can change for a species (Vander Zanden *et al.* 1999; Tewfik *et al.* 2016). Our goal was simply to develop a general assessment of a change in fish assemblages, and we considered the trophic position provided by Fishbase (Froese & Pauly 2015) as a reasonable proxy to evaluate if fish get more predatory over time in reservoirs. Follow up studies using more direct approaches (*e.g.,* stable isotopes) would be worthwhile to investigate this observation more fully. Finally, what constitutes an adequate reference ecosystem for a reservoir, and the potential differences in ecosystem sizes among studies and biomes need consideration. In cross-sectional datasets, the unregulated sites for boreal ecosystems were all lakes whereas unregulated sites were mainly rivers and streams in temperate (5% lakes, 95% rivers or streams) and tropical ecosystems (11% lakes, 89% rivers or streams). We need appropriate reference ecosystems to control for stochasticity and climatic events, but comparing reservoirs to only reference lakes or only rivers might be inadequate because reservoirs are neither a lake nor a river. Only one study compared reservoir fish communities with those in rivers and lakes in temperate systems and found that reservoir communities were more similar to lake vs. river communities (Irz *et al.* 2006). Differences in ecosystem size between reservoirs and lakes is another plausible explanation for the trophic position and diversity results presented herein, as there is certainly a well-established body of literature showing that these metrics scale with ecosystem size (Post *et al.* 2000). Similarly, geographic location is known to influence fish richness in lakes (Matuszek & Beggs 1988; Samarasin *et al.* 2014). We clearly see that fish diversity metrics are higher in tropical sites (even before impoundment), which is consistent with the expected trend. We believe that part of the effect of ecosystem size is reflected in the regression tree that explores what variables might explain differences in RE among studies, where we found that catchment area was the strongest predictor. Awareness of the potential effects associated with ecosystem size is particularly relevant when investigators are comparing reservoirs to natural lakes. Boreal reservoirs used in this synthesis were on average 158 times bigger than adjacent reference lakes. Therefore, richness should be higher in reservoir relative to adjacent reference lakes just based on their respective size. Empirical and experimental studies going forward would be well advised to take these factors into consideration.

### Concluding remarks

Based on an analysis of 147 longitudinal and 37 cross-sectional studies from a total of 67 references, we present a comprehensive synthesis that quantitatively evaluates the effects of impoundment on fish biodiversity and species assemblages across three globally-dominant biomes. Four major insights emerge from our synthesis. First, predictions regarding the impact of dams on fish communities require a regional perspective. Tropical regions were more affected and characterized by stronger changes in richness and diversity, and marked increases in non-native species following impoundment. In contrast, lower amplitude changes were observed in temperate and boreal reservoirs. Second, the full extent of fish communities’ dynamics in tropical regions remains to be determined as time series here are short. Tropical reservoirs are young and novel ecosystems, and are still in the non-equilibrium phase. Third, a lack of change in richness does not mean no change in native species richness. We observed a sharp increase in non-native species in the tropics that was not observed in boreal ecosystems, and this effect masked changes in the whole fish assemblage. Finally, changes in fish assemblages is a common feature across regulated ecosystems. We detected a global increase in the trophic level position and a general decrease in the percentage of rheophilic species. Collectively, we conclude that the changes in fish assemblages and diversity detected in reservoirs could potentially impact the stability of the food web, the productivity of these ecosystems, the sustainability of artisanal fisheries, and the function and ecosystems services (Hoeinghaus *et al.* 2009; Toussaint *et al.* 2016). In light of this global quantitative synthesis, hydropower may be part of the solution to decarbonize our global economy but will come at substantially higher ecological cost to the tropics (Ziv *et al.* 2012; Winemiller *et al.* 2016; Pelicice *et al.* 2017). When planning hydropower development, strategic and transboundary actions should be taken to protect, conserve and restore fish biodiversity, particularly in the sensitive regions like the tropics.

## Acknowledgements

This work was supported by a MITACS Elevate scholarship to KT with Hydro-Québec as an industrial partner. IGE also acknowledges support from the Canada Research Chairs program. We thank J. Caumartin (Hydro-Québec), J.-C., Guay (Hydro-Québec), P. Johnston (Hydro-Québec), C. Solomon (Cary Institute), C. Martin (UQTR), A. Tremblay (Hydro-Québec) and B. Vanier (Hydro-Québec) for helpful suggestions and comments on an early draft of this manuscript.

